# Predicting genetic biodiversity in salamanders using geographic, climatic, and life history traits

**DOI:** 10.1101/2024.02.16.580580

**Authors:** Danielle J. Parsons, Abigail E. Green, Bryan C. Carstens, Tara A. Pelletier

## Abstract

The geographic distribution of genetic variation within a species reveals information about its evolutionary history, including responses to historical climate change and dispersal ability across various habitat types. We combine genetic data from salamander species with geographic, climatic, and life history data collected from open-source online repositories to develop a machine learning model designed to identify the traits that are most predictive of unrecognized genetic lineages. We find evidence of hidden diversity distributed throughout the clade Caudata that is largely the result of variation in climatic variables. We highlight some of the difficulties in using machine-learning models on open-source data that are often messy and potentially taxonomically and geographically biased.

## Introduction

Documenting biodiversity is an important first step in understanding both ecological and evolutionary processes (Gadelha *et al.*, 2021), particularly the functional roles that act to connect processes functioning at both shallow and deep time scales (Guralnick & Hill, 2009). Notably, any such documentation of biodiversity implicitly assumes that the units (e.g., species) are comparable across different geographic regions. Given that a Linnean shortfall (i.e., the ratio of recognized to unrecognized species (Whittaker *et al.*, 2005)) exists in most clades and may be substantial across Eukaryota (Mora *et al.*, 2011), it is not clear that this assumption is reasonable. An alternative approach is to utilize evolutionary significant units (Moritz, 1994), or genetic lineages, in place of species in broad analyses of biodiversity (e.g., (Mable, 2019)). This may be particularly useful in clades with relatively high degrees of morphological and ecological conservatism. One such clade is Caudata (i.e., salamanders and newts), which exhibits high frequencies of cryptic species (e.g., (Jockusch *et al.*, 2012; Camp & Wooten, 2016; Bernardes *et al.*, 2020)).

Identifying genetic lineages in Caudata can have important conservation implications. For example, Mead *et al.* (2005) discovered a new species of western *Plethodon* salamander that was originally thought to be either *P. elongatus* or *P. stormi* (Mead *et al.*, 2005). All three of these species are listed on the IUCN Red List as either near threatened (*P. elongatus*), vulnerable (*P. asupak*), or endangered (*P. stormi*). More recently, Parra Olea *et al*. (2020) discovered five cryptic lineages in *Chiropterotriton* from Mexico, several of which are threatened due to their restricted ranges (Parra Olea *et al.*, 2020). Species with small ranges and/or limited dispersal capabilities can be harder to protect because their distributions often do not fall within protected areas (Nauman & Olson, 2008) and small ranges are often used as a factor in assigning conservation priorities (Hortal *et al.*, 2015). Therefore, it is important to identify these lineages, as they could easily go unnoticed and unprotected. Many other species of salamander that would have otherwise gone unnoticed and have been recognized using molecular data have small ranges and likely need protection (Steffen *et al.*, 2014; Nishikawa & Matsui, 2014; Min *et al.*, 2016; Kuchta *et al.*, 2018; Okamiya *et al.*, 2018). The presence of cryptic diversity has been recently highlighted as a key component of undescribed biodiversity that requires greater attention (Bickford *et al.*, 2007; Pfenninger & Schwenk, 2007).

Efforts to conserve undescribed genetic diversity can be facilitated using computational methods that identify genetic lineages representing potentially hidden diversity in need of further investigation. The use of data science techniques has allowed biodiversity studies to expand their geographic and taxonomic focus to explore broader patterns of evolution, which can be difficult to assess using traditional meta-analysis methods (Lyman & Edwards, 2022). Macrogenetics, a relatively new field that merges biodiversity data with genetic data (Blanchet *et al.*, 2017; Leigh *et al.*, 2021), has been used to explore how human impacts influence levels of intraspecific genetic diversity (Miraldo *et al.*, 2016; Millette *et al.*, 2020), to study past and future climate refugia (Carstens *et al.*, 2018; Baranzelli *et al.*, 2022), and to quantify latitudinal biodiversity gradients (Gratton *et al.*, 2017; Pelletier & Carstens, 2018; Barrow *et al.*, 2021; Fonseca *et al.* 2023). Macrogenetic methods, particularly in combination with predictive modeling, can be used to inform conservation policies by identifying species, taxonomic groups, or geographic areas in need of further investigation (Pelletier *et al.*, 2018; Raposo *et al.*, 2021). Recently, such analyses have expanded to taxonomic work.

Parsons *et al*. (2022) analyzed mitochondrial DNA sequences from over 4000 species of mammals, representing roughly 66% of currently described species, and found that mammal diversity is largely under-described using molecular species delimitation methods on publicly available barcode data. This is useful for several reasons. A comprehensive list of genetic lineages that may represent species now exists that can help focus taxonomic efforts. Parsons *et al.* (2022) also found that taxa with small bodies, and large geographic distributions with variation in precipitation and isothermality, were more likely to contain cryptic diversity. While some of this might seem obvious (morphological differences are harder to observe in small-bodied animals and these animals may be harder to find), it does allow researchers to document characteristics of species, higher taxonomic groups, or even geographic regions that contribute to diversification and therefore biodiversity patterns. When done in disparate taxonomic groups (e.g., vertebrates, invertebrates, plants, and fungi) and at different levels (e.g., Class, Order, Family) this furthers our understanding of core evolutionary processes.

A similar approach was taken in birds. Using a tree-based molecular species delimitation method, Smith *et al.* (2018) found that latitude explained variation in phylogeographic breaks, while other traits pertaining to habitat and life history explained very little. In this case, phylogeographic structure was higher in the tropics. Conversely, in other organisms, isolation-by-distance within species is often higher at higher latitudes (multiple taxonomic groups: Pelletier & Carstens, 2018; amphibians: Amador **et al.*,* 2023). Further, genetic variation within amphibians was best explained by range size and elevation, rather than latitude, in the neotropics (Amador **et al.*,* 2023), while latitude was an important predictor of genetic diversity in the nearctic (Barrow *et al*., 2021). This suggests that differences exist in how genetic variation is distributed within species depending on which taxonomic groups are being examined, and at what spatial scale.

In order to expand these approaches, we conducted a computational assessment of genetic lineages in roughly 100 salamander species using the *phylogatR* database (Pelletier *et al.*, 2022). *PhylogatR* aggregates DNA sequence data from both GenBank and BOLD into sequence alignments, providing associated GBIF occurrence records (i.e., GPS coordinates) for each sequence. There are over 700 described species of salamanders belonging to nine families (Bánki, O. *et al.*, 2022), most located in the northern hemisphere. While salamanders contain a wide variety of life history strategies and habitats, they are likely to have high levels of cryptic diversity due to their moisture requirements and similar body forms. However, their eco-evolutionary processes can vary from species to species and sometimes oppose our expectations (Pelletier *et al.*, 2011, 2015; Pelletier & Carstens, 2016; Jones and Weisrock 2018; Pyron **et al.*,* 2020; Dufresnes *et al.*, 2021). We follow methods from Parsons *et al.* (2022) and use molecular species delimitation methods to estimate the number of genetic lineages present in previously collected data that is both openly available and easily tractable. We then use a predictive modeling approach to determine whether any variables pertaining to geography, the environment, or life history traits contribute to the presence of genetic lineages within species. We also discuss some of the difficulties in using open-source data that are often messy and potentially taxonomically and geographically biased.

## Materials and Methods

### Collection of genetic and geographic data

We downloaded all available data from the *phylogatR* database (https://phylogatr.org/) using the search term ‘Caudata’ on 2/4/22. The uncleaned data represented four families, 93 different species, and 14 loci with a total of 3768 DNA sequences. To begin cleaning the data, we calculated nucleotide diversity (pi) values for each locus in every species and found outliers by setting lower and upper bounds of 2.5% (0) and 97.5% (0.2193634) respectively. For each of the four outliers and two species with missing pi values, we opened the DNA sequence file in Mesquite v3.7 (Maddison & Maddison, 2021) and removed any extremely short or non-overlapping sequences (Data S1). Additionally, we discovered a typo for the species *Batrachuperus karlschmidti* causing there to be two different species folders for the same species. Both the sequence and occurrence files were merged for the species and the sequence files were realigned to correct the error. Two species complexes were present in the dataset, and these were kept named as downloaded: *Triturus cristatus x dobrogicus macrosomus* and *Ambystoma laterale jeffersonianum* complex.

Species alignments from the download for both the mitochondrial genes Cytochrome oxidase I (*COI*) and Cytochrome b (*cytb*) were merged for all salamander species and aligned using MAFFT v7.5 (Katoh & Standley, 2013) with the default settings and including the – adjustdirection command to account for reverse complement sequences. We visually inspected alignment files for both genes and removed all short sequences, which we classified as those missing 50% or more of the second half of the sequence. Twenty-one sequences were removed from the *COI* alignment and 99 were removed from the *cytb* alignment, leaving totals of 768 and 908 sequences for *COI* and *cytb*, respectively. The sequences for seven species were completely removed from further analysis due to their short length (missing 50% or more of the second half of the sequences). In total, eighty-three species remained with an average of approximately 20 sequences per nominal species (see Data S2 for a list of identifiers corresponding to the sequences used in this study).

### Species delimitation

We used three methods of species delimitation to determine the number of genetic lineages present in our samples. The GMYC is a tree-based method that takes a phylogenetic tree as input and finds a point in the tree where branching changes from within to between species (Pons *et al.*, 2006). The ABGD (Puillandre *et al.*, 2012) and ASAP (Puillandre *et al.*, 2021) methods are distance-based delimitation methods that use pairwise genetic distances to establish the threshold between intra- and inter-species divergence. Because each method is based on a specific set of assumptions, it is best to use multiple methods and compare their results in order to achieve a more accurate delimitation (Carstens *et al.*, 2013). By looking for concordance across methods, we can increase our confidence in the identified lineage boundaries and minimize the potential impact of bias introduced by any single method. While we report delimitation results from the genes *COI* and *cytb* for all methods, we used a consensus of delimitation results (among methods and loci) for assessing the role of geography, the environment, and life history traits in predicting salamander genetic diversity.

To estimate a species tree for input into the GMYC, we used BEAST v2.5.1 (Bouckaert *et al.*, 2019). We used the default parameters except for conducting 100,000,000 million generations, sampling every 5,000, and setting the model of sequence evolution to GTR+I+G (Abadi *et al.*, 2019). The log files were checked by eye using Tracer v1.7.2 (Rambaut *et al.*, 2018). ESS values were all over 1000 for both *cytb* and *COI*. We removed 10% as burnin and retained the maximum clade credibility tree using TreeAnnotator. After checking that the tree was binary and ultrametric, we used the R package *splits* (Ezard *et al.*, 2009) to conduct GMYC analyses. In each case we used the single threshold model and all other default settings. We conducted both ABGD and ASAP delimitation analyses via their web portals (https://bioinfo.mnhn.fr/abi/public/abgd/abgdweb.html and https://bioinfo.mnhn.fr/abi/public/asap/asapweb.html, respectively) using the default parameter settings.

### Predictor variables

A variety of predictor variables were collected, including geographic and environmental values derived from georeferenced locality data (see Data S3). In addition, three life history traits were available from AmphiBIO, a global database for amphibian ecological traits (Oliveira *et al.*, 2017), for most of the species in our study: reproductive strategy (direct developing, larval phase), habitat (terrestrial, fossorial, aquatic, or some combination of these), and body size (total length). To supplement this dataset and fill in any missing trait values, we used AmphibiaWeb (AmphibiaWeb, 2023) and other online sources (Data S4).

To extract species specific data related to its environmental distribution, we utilized 42 GIS data layers (see Data S4 for data layer details), including all 19 BIOCLIM layers from the CHELSA database (Karger *et al.*, 2017; Karger, Dirk Nikolaus *et al.*, 2021) at 1 km resolution, elevation (Aster global digital elevation model version 2, 2011), population density (Socioeconomic Data And Applications Center (SEDAC) Gridded Populations of the World (GPW), 2016), terrestrial habitat heterogeneity (Tuanmu & Jetz, 2015), gross domestic product (World Bank Development Economics Research Group (DECRG) Gross Domestic Product, 2010), global land cover classification (European Space Agency, 2009), global river classification (Ouellet Dallaire *et al.*, 2019), disaster risk (Peduzzi, 2019), anthropogenic biome (Ellis *et al.*, 2010), and various indicators of seasonal growth (Karger *et al.*, 2017; Karger, Dirk Nikolaus *et al.*, 2021). We utilized the R packages ‘raster’ (, 2016), ‘rgdal’ (, 2017), ‘geosphere’ (,2016), and ‘plyr’ (Wickham, 2011) to extract species specific information from each layer using geographic occurrence records obtained from *phylogatR*. To represent the environmental variation within the occupied range of each species, we extracted the value of each environmental layer for each GPS coordinate associated with each species. We then took the mean and standard deviation for each environmental variable. To obtain species specific data related to geographic distribution we extracted the minimum, maximum, mean, and length of latitude and longitude from the GPS points of each species.

We used the R package ‘mice’ (Buuren & Groothuis-Oudshoorn, 2011) to impute trait values missing from our dataset (see Figure S1 for distribution of missing data and specific trait values imputed). The imputation method **’**pmm**’** was used for all numeric variables and **’**polyreg**’** was used for categorical variables (i.e., reproductive strategy and habitat). We ran the imputation 15 times (Figure S2) and then pooled the iterations to generate the final imputed values. The final database containing all trait values (both imputed and original) is available in Data S4.

### Predictive modeling

We used the R package ‘caret’ (Kuhn, 2008) to generate a random forest classification model (Breiman, 2001) based on our previously generated database of predictor variables and a consensus of our species delimitation results. Two separate sets of consensus models were generated to assess the role of geography, environment, and life history traits on the presence of hidden diversity (Figure 1A). The first model (*all agree*) represents a strict consensus of delimitation results from species in which results from all methods of species delimitation agree (Figure 1B). Any species with conflicting delimitation results were excluded from analysis. The second model (*majority rules*) represents a majority rule consensus in which species are assigned to a response category based on relative support of delimitation results (Figure 1C). For each model, we used 70% of the data to train the model and the remaining 30% was set aside as a test set. Models were generated using 10-fold cross validation with five repeats to tune the parameter ‘mtry’, the number of variables randomly sampled at each split, and optimize the area under the receiver operating characteristic curve, ROC. After training, we extracted the variable importance measures mean decrease accuracy (MDA) and Gini impurity (Gini) from the final models. We then used the final models on the test set data to evaluate model performance. Model performance was evaluated across a variety of metrics including model accuracy, which reflects how well the predicted classifications agree with the observed classifications, and both positive and negative predictive value, which indicate the how the model performs on observations from each class. Additionally, we calculated the no information rate (NIR), the proportion of observations that fall into the majority class, and the p-value [Accuracy>NIR], to test for model significance. The top important predictor variables from our best model were compared using a Kruskal-Wallis test to determine if these variables are significantly different between species that do or do not contain hidden diversity.

**Figure 1.**
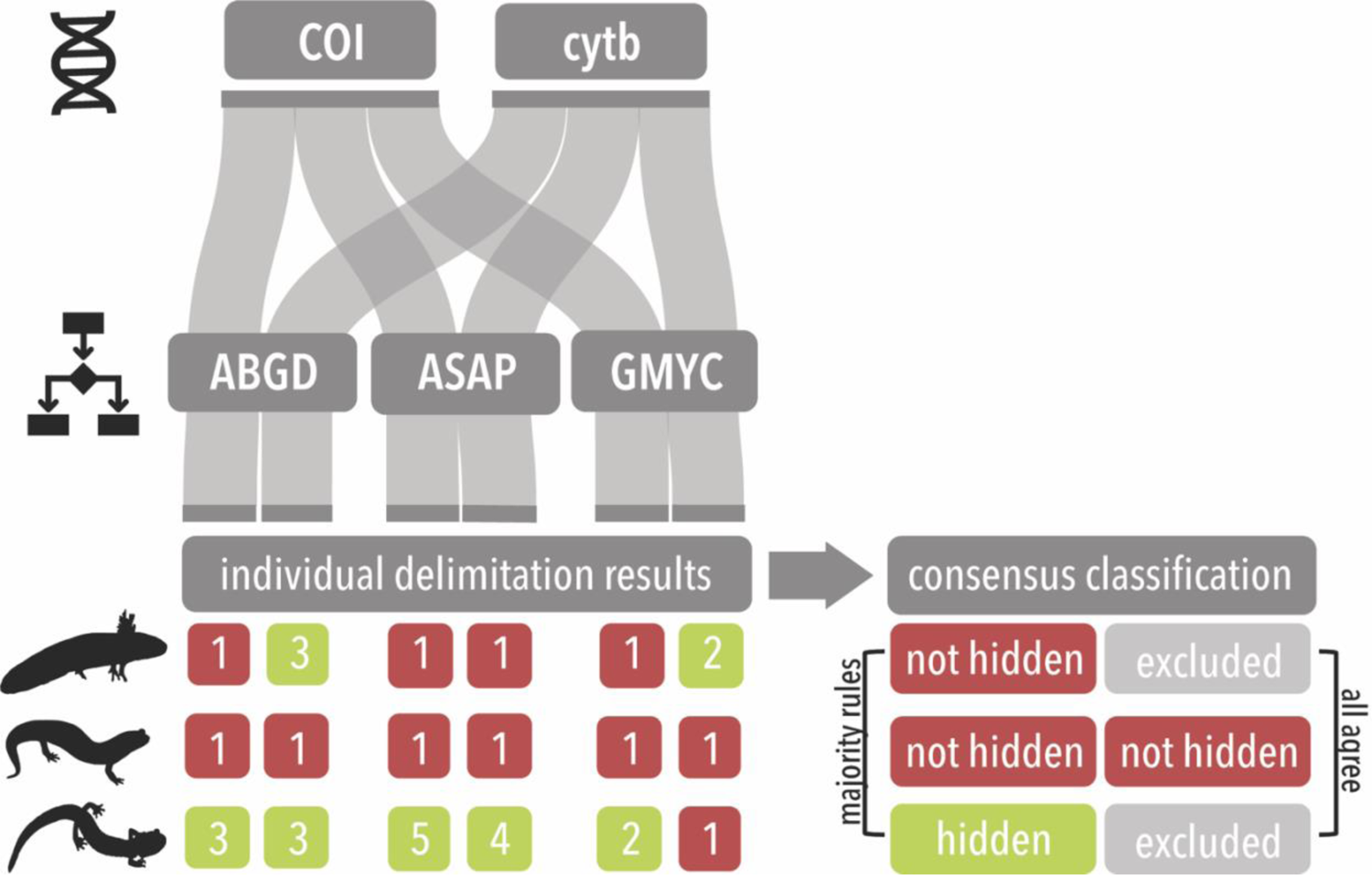
Consensus classification of species delimitation results. A, Flowchart describing the process of generating a consensus of delimitation results (among different methods and loci). B, C, Pipeline for classifying nominal species as either containing or not containing hidden diversity in each consensus analysis (*all agree* and *majority rules*, respectively).

## Results

### Genetic and geographic dataset

Our final dataset consisted of 1676 DNA barcoding sequences (Figure 2). Of these, 768 sequences were from the Cytochrome oxidase I gene (*COI*), and 908 sequences were from the Cytochrome b gene (*cytb*). These sequences were derived from 83 nominal species of salamanders, which were distributed among 26 distinct genera occurring across the globe. The dataset contained 13 species with sequences from the gene *cytb*. Comparatively, *COI* exhibited notably broader taxonomic coverage, with 77 nominal species represented. Out of the 83 species analyzed, only seven were shared between *COI* and *cytb*. Of the remaining 76 species, 70 were unique to *COI* and six were unique to *cytb*. To supplement the genetic data collected, a total of 1765 georeferenced occurrence records from *phylogatR* were utilized to collect a combination of geographic, environmental, and life history trait values for each nominal species present in the dataset.

**Figure 2.**
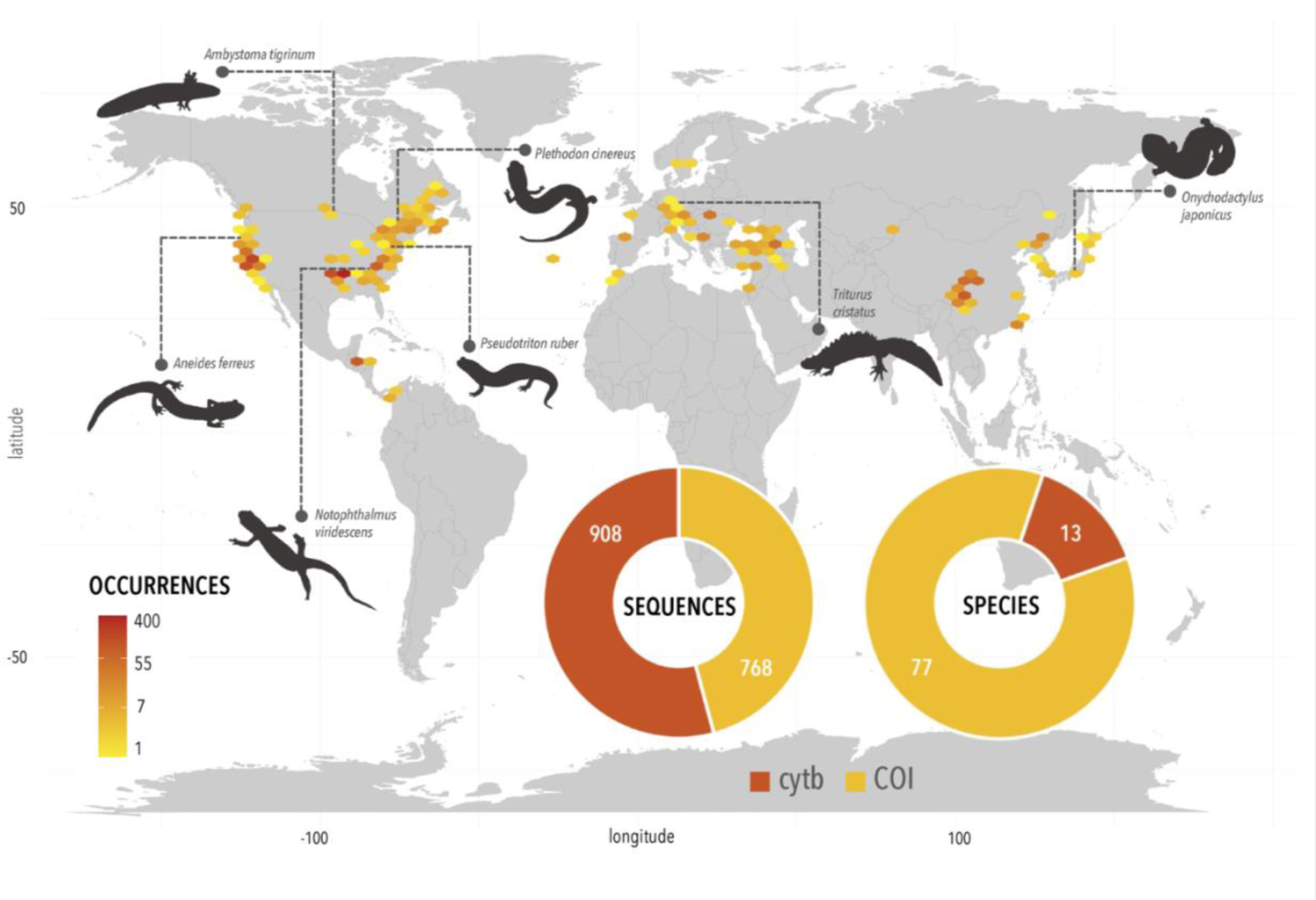
Geographic spread of salamander data. Map shows geographic distribution of salamander occurrences pulled from *phylogatR* (Pelletier *et al.*, 2022) and used in these analyses. Pie charts show the total number of *cytb* and *COI* sequences used (left) and the number of species represented by those *cytb* and *COI* sequences (right). Salamander figures in black were obtained from Phylopic (M. Keesey) and are licensed under public domain.

### Species delimitation and consensus assignment

Species delimitation results were generated by analyzing *COI* and *cytb* sequences from each nominal species under three different delimitation methods, ABGD, ASAP, and GMYC. We classified each nominal species as either containing genetic lineages or not containing genetic lineages based on the number of genetic groups predicted by each delimitation analysis. While taxonomic overlap between *COI* and *cytb* was narrow, delimitation results for species shared by both loci were mostly congruent with respect to species classification. Of the seven species with sequences from both genes, only two species produced conflicting results regarding the presence of genetic lineages within a specific taxon based on loci. Delimitation results across different methods showed slightly less agreement. Classifications resulting from the GMYC and ASAP methods were similar across species. These methods, on average, resulted in slightly fewer predicted species per nominal species than the ABGD method (see Figure 3 for predicted species numbers).

**Figure 3.**
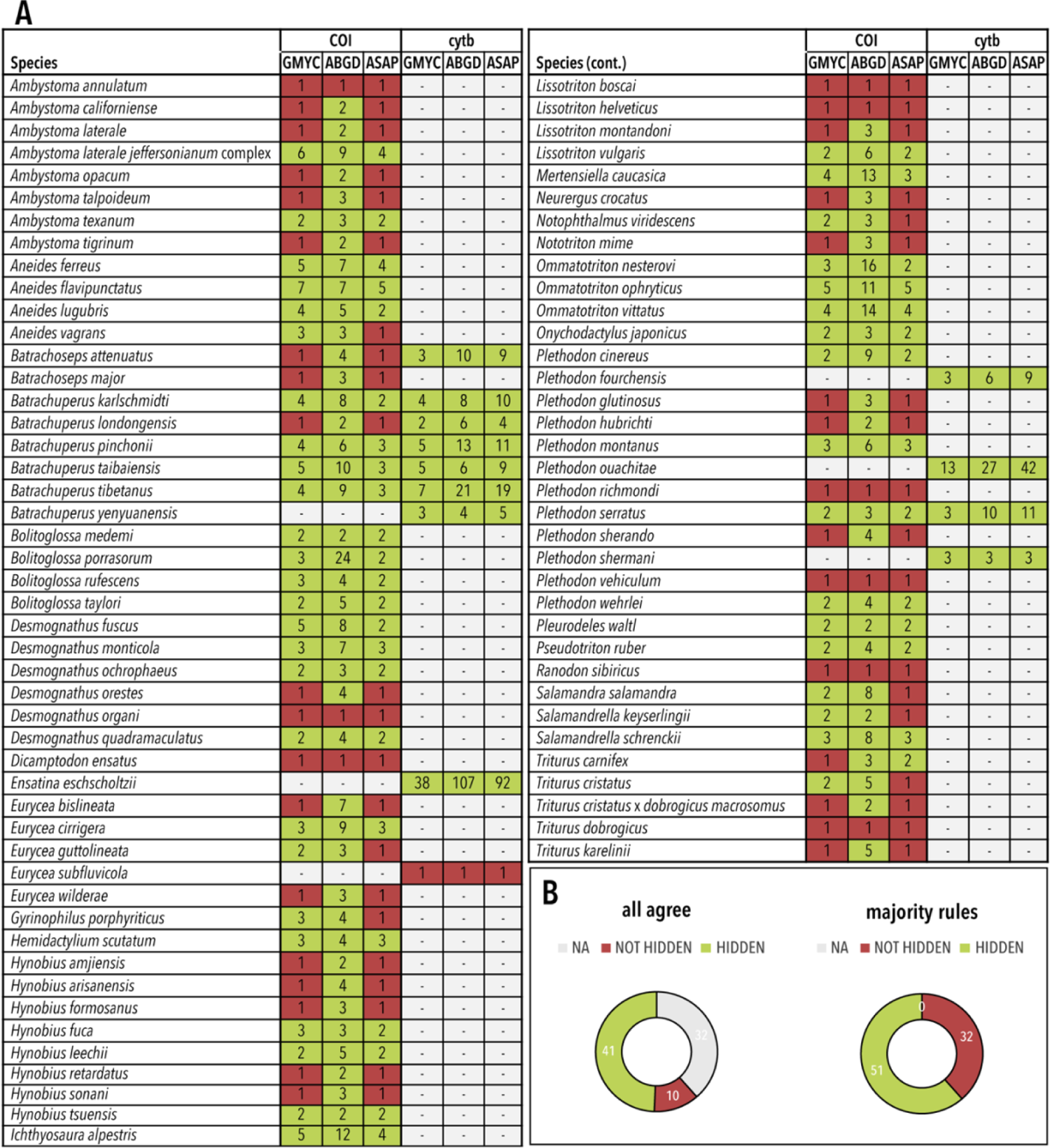
Species delimitation results. A, Graphs show the results of ABGD, ASAP, and GMYC species delimitation analyses of the genes *cytb* and *COI* for each nominal species. Numbers represent the predicted genetic lineages from each analysis. Results highlighted in red indicate no hidden genetic lineages were predicted (i.e., number of genetic lineages = 1). Results highlighted in green indicate hidden genetic lineages were predicted (i.e., number of genetic lineages > 1). Grey highlighting indicates that specific analysis was not performed due to a lack of data. B, Pie charts display the number of nominal species classified as either containing or not containing hidden diversity in each consensus analysis (i.e., *all agree* and *majority rules*).

To account for this variation in our final predictive models, we generated two consensus classifications to evaluate concordance between delimitation results from different methods and loci. The results of our consensus models indicate that roughly 2/3rds of the nominal salamander species used in this analysis are likely to contain genetic lineages that may be unexplored diversity. The strictest of these classifications produced a consensus model (*all agree*) consisting of 51 total species, 41 of which were classified as containing hidden diversity and 10 of which were classified as not containing hidden diversity. The remaining consensus model (*majority rules*) consisted of 83 total species, of which 51 were classified as containing genetic lineages and 32 were not (Figure 3).

### Predictive modeling

For our *majority rules* and *all agree* consensus classifications, we developed random forest classification models using all available predictor data. To assess potential correlation between variables in our dataset we used the R package ‘corrplot’ (Taiyun Wei & Viliam Simko, 2021) to generate a correlation matrix of our predictor variables (Figure S3). Due to the presence of strong correlations between several of the geographic and environmental variables in our dataset we performed multiple random forest models with progressive sets of correlated variables removed at different cutoff values (i.e., |correlation coefficient| > 0.75; 0.85; 0.9). The results of these random forest models are presented below (Table 1).

**Table 1.** Results of *majority rules* consensus predictive models. Model metrics for each random forest predictive model generated using the *majority rules* consensus classifications are shown.

| Majority Rules<br>Models | Original | \|Correlation\| > 0.75 | \|Correlation\| > 0.85 | \|Correlation\| > 0.90 |
| --- | --- | --- | --- | --- |
| Accuracy | 0.75 | 0.75 | 0.75 | 0.8333 |
| Accuracy (95% CI) | (0.5329, 0.9023) | (0.5329, 0.9023) | (0.5329, 0.9023) | (0.6262, 0.9526) |
| No Information Rate | 0.625 | 0.625 | 0.625 | 0.625 |
| Pos Pred Value | 0.7368a | 0.8 | 0.7647 | 0.7895 |
| Neg Pred Value | 0.8 | 0.6667 | 0.7143 | 1 |
| P-Value [Acc > NIR] | 0.1453 | 0.1453 | 0.1453 | 0.02435 |

All random forest models were found to have high predictive accuracy, with the *majority rules* and *all agree* models achieving accuracies of 75-85% and 87-93%, respectively, in identifying nominal species likely to contain hidden diversity. Although these results may initially seem to suggest that all our models are able to make meaningful predictions, further examination of additional model evaluation metrics reveals potential overfitting and inflation of predictive power. For example, despite the high accuracy of the models, the 95% confidence intervals for these values are broad with an average length of nearly 40% for most of the models (Tables 1 and 2). Additionally, the no information rates (NIRs), a measure of prediction significance based on the underlying dataset that needs to be exceeded in order for model results to be significant, are particularly high for the *all agree* consensus models, where the class frequencies are more skewed towards species predicted to harbor hidden diversity. The high NIR values combined with wide confidence intervals result in a p-value [Accuracy > NIR] greater than 0.05 in all models, except for the *majority rules* consensus using a correlation cutoff of 0.90. While all our models show high accuracy, when the additional model evaluation metrics are considered only one has strong predictive power. Therefore, we only used the *majority rules* consensus using a correlation cutoff of 0.90 for interpreting variable importance of our data.

**Table 2.**
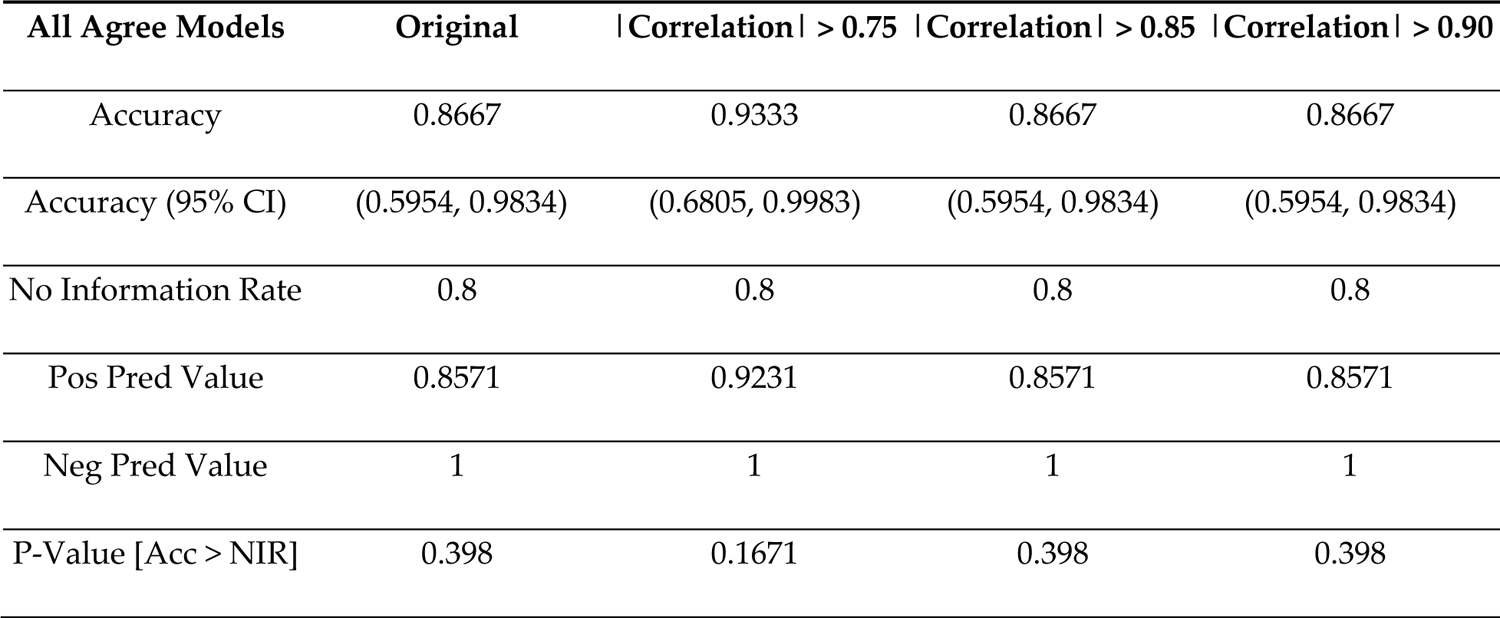
Results of *all agree* consensus predictive models. Model metrics for each random forest predictive model generated using the *all agree* consensus classifications are shown.

### Evaluation of variable importance

We extracted variable importance measurements from each predictive model using the variable importance metrics MDA and Gini. While there was some overlap of top predictors between different models (Figure 4; Figure S4), no specific predictors were consistently predicted to be of significantly higher importance than other predictors in the model. Instead, importance was split across numerous predictors that were found to be unstable between models. This instability supports previous indications that many of the predictive models are likely prone to overfitting. Despite the lack of a strong set of standout predictors across models, one pattern does emerge that is applicable to the species in our dataset. Of the top ten most important predictors in each model, approximately 85% are measurements of standard deviation (vs. measurements of mean values or life history traits) (Data S5). This is supported by further examination of our one model that was able to predict significantly better than random, the *majority rules* consensus with a correlation coefficient cutoff of 0.90, in which the top five most important predictors are measurements of standard deviation. Significance testing indicates that species identified as containing hidden genetic lineages often have ranges characterized by a larger variance in annual and seasonal precipitation, isothermality, and net primary productivity than species not identified as harboring hidden genetic lineages (Figure 5).

**Figure 4.**
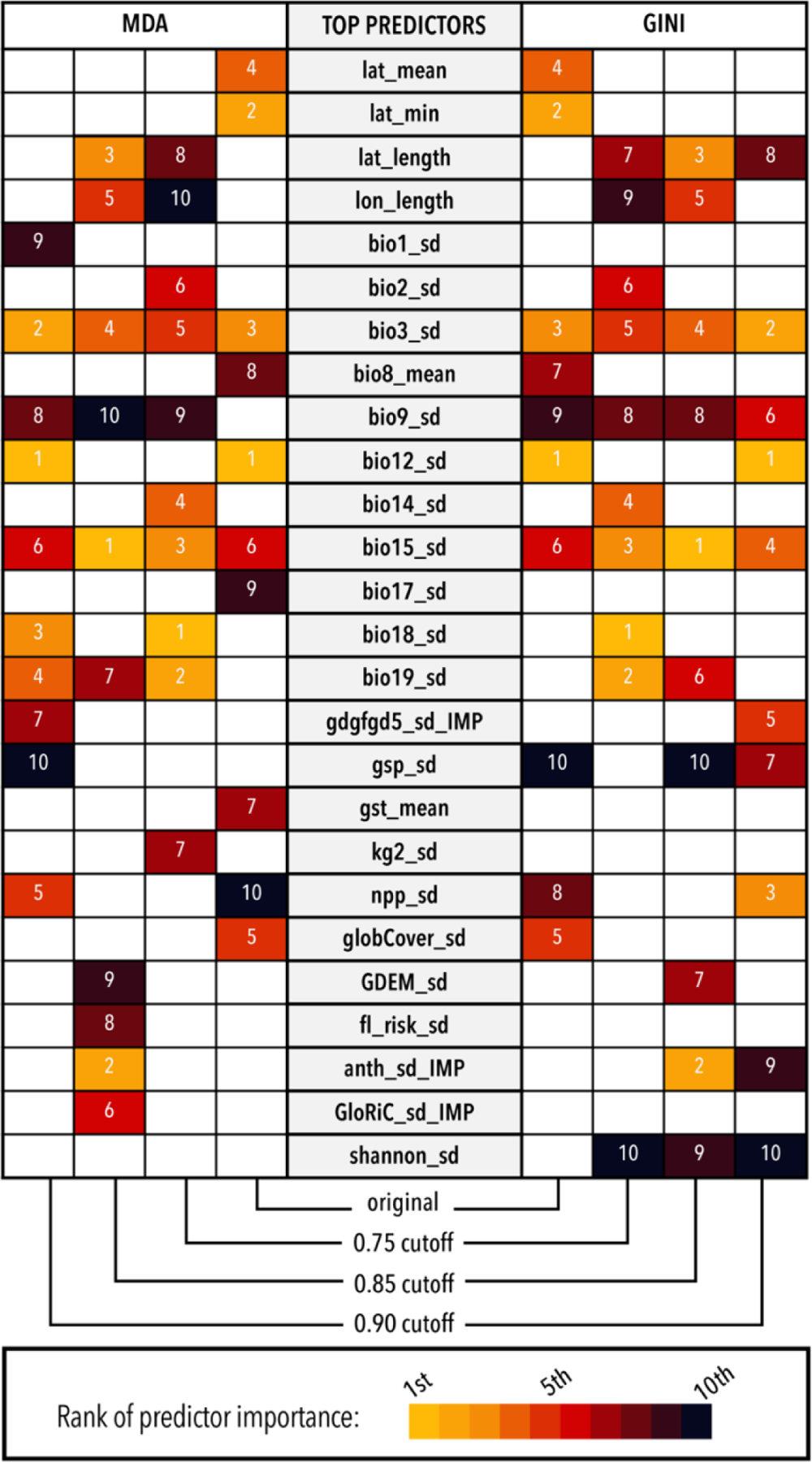
Variable importance for predictive models generated using the *majority rules* consensus. Variables ranked among the top ten most important variables (based on MDA and Gini) from the predictive model generated at different correlation cut-offs are included.

**Figure 5.**
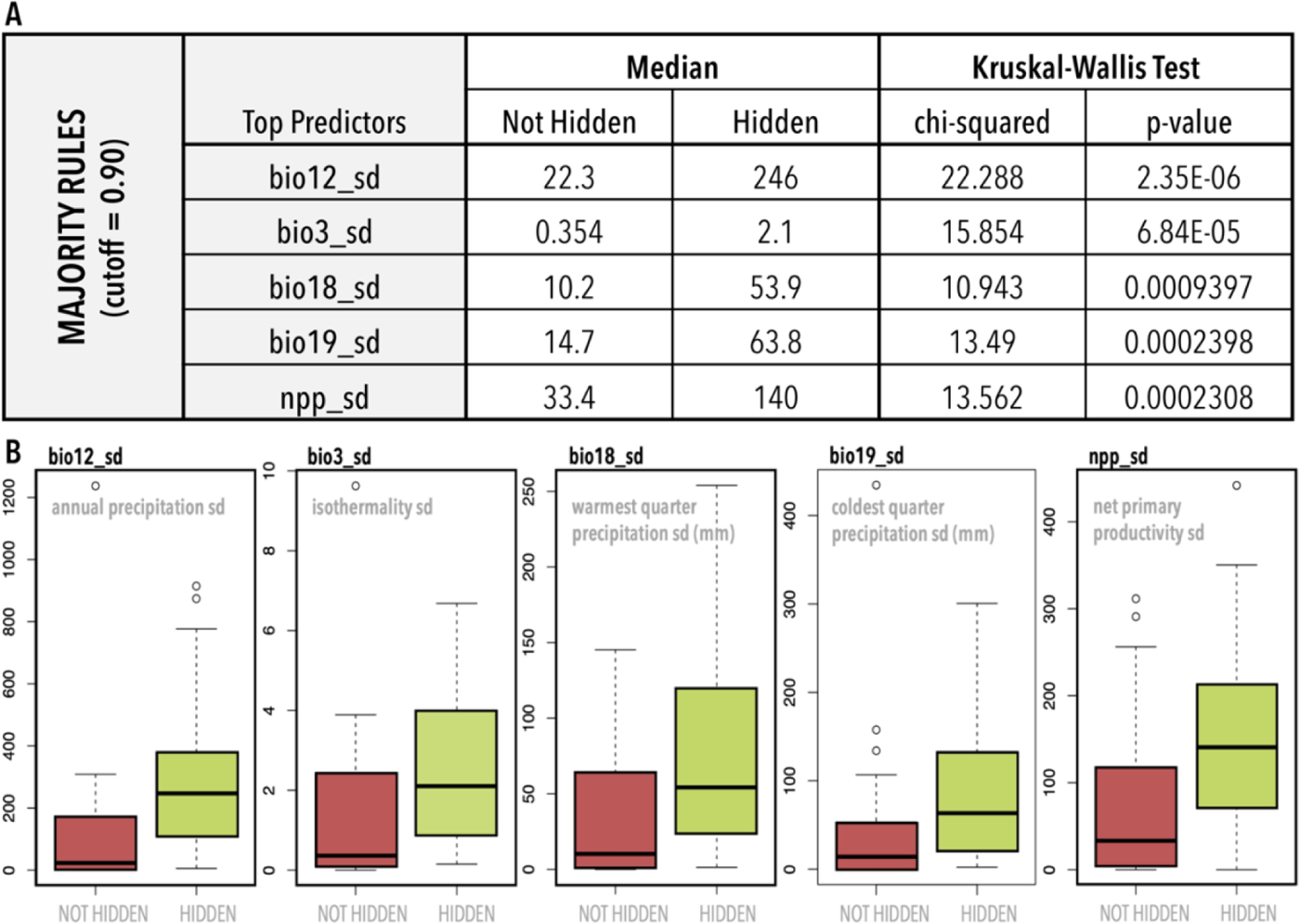
Difference in hidden vs not hidden trait values. A, Results of Kruskal-Wallis significance test on the top five most important predictors of the best model (*majority rules* – correlation cutoff 0.90). B, Corresponding boxplots for said predictors show a significant difference in the range of trait values between hidden and non-hidden genetic lineages.

## Discussion

When identifying genetic lineages or delimiting species, it is important to recognize that species concepts are complex and often differ based on various factors, such as geographic location, reproductive isolating mechanisms, genetic markers, and taxonomic practices. Therefore, it is essential to approach species delimitations with caution and to recognize that they represent a hypothesis or starting point rather than a definitive answer (Hillis, 2019). In addition, while mitochondrial data can be suitable for preliminary assessments of species diversity (Gostel & Kress, 2022), these assessments should be considered in tandem with other species information and relevant data when describing species boundaries. However, with recent advances in technology rapidly increasing the quantity of publicly accessible genetic and geographic datasets, these data offer a cost effective and efficient way to explore large-scale patterns and predictors of intraspecific genetic variation (e.g., Miraldo *et al.*, 2016; Pelletier & Carstens, 2018; Yiming *et al.*, 2021).

Our results suggest that there are genetic lineages that may warrant further investigation distributed within Caudata. Adequately documenting biodiversity, both at the species and population level, is a first step in understanding the eco-evolutionary processes generating this diversity. However, in most clades, the Linnean shortfall is likely to influence broad scale patterns detected using macrogenetic approaches (Hortal *et al.*, 2015), making it essential to consider how the taxonomic designations used to inform these approaches influence the patterns detected. This is particularly important when dealing with clades suspected of harboring high levels of cryptic diversity. For example, Miraldo *et al*. (2016) generated the first global map of genetic diversity within species of mammals and amphibians. One of their main conclusions was that amphibians displayed lower levels of genetic variation in areas with higher human impact. Similarly, in amphibians, several recent studies have found within species genetic diversity to be lower in temperate regions in species with smaller ranges and at higher elevations (Barrow *et al*., 2020; Amador 2023). The methods used to detect these patterns are based on current taxonomic knowledge, and as such, rely on the assumption that the species designations used are accurate. However, if species descriptions inaccurately reflect biological diversity, nominal species that contain cryptic species will display higher levels of genetic diversity, while not reflecting true within species variation, potentially skewing our interpretation of any patterns that result.

### Evaluating support for identified genetic lineages

While our delimitation of genetic lineages are a starting point, or hypothesis generation step, for evaluating a species in nature where complex processes, such as hybrid zones, and adequate sampling must be considered (Hillis, 2019), we believe these computational approaches are useful for targeting species in further need of examination. We conducted a literature search to explore whether the nominal species in our dataset have been previously explored from a species delimitation approach. We used the online American Museum of Natural History taxonomic and nomenclatural database, Amphibian Species of the World (Darrel, 2024), to evaluate current taxonomic research in each nominal species of salamander predicted to contain hidden diversity in our consensus model. Species in which we were able to identify research-based support for the potential of undescribed diversity were recorded, along with the related articles in which the diversity was described as well as the type of data used (see Data S7). Nearly 70% of species the majority rules consensus suggests harbor hidden lineages contain results that also support the potential splitting of species into separate lineages. Out of these about 38% were explored using mt DNA only, 10% with nuclear DNA only, 35% using a combination of both nuclear and mt DNA and 17% using mt DNA, nuclear DNA and morphology. Just under 10% of the species display a complex history of hybridization, making delimitations difficult, a situation not uncommon in salamanders (Denton *et al*., 2018; Pyron *et al*., 2020). We were unable to find results for roughly 25% of our species data. We encountered 5 species in which the results of previous delimitation work was either unclear or considered highly contested (e.g., *Ichthyosaura alpestris*, *Batrachuperus karlschmidti*, *Batrachuperus taibaiensis,* and *Salamandrella schrenckii*). Taxonomy is dynamic field (Raposo **et al.*,* 2020) and given our search, it can be difficult to use current open-source data relying solely on species names. However, the current literature largely supports the delimitation results found here and suggests a number of species in further need of investigation (see citations in Data S7, formal name changes, and an ability to update current open-source databases to reflect these changes). Additionally, even though there are limitations to using current open-source data that might not keep up to date with current taxonomy, we can still determine what factors might predict the presence of hard-to-find species.

### Significant Predictors of Diversity

Significance testing of the most important predictors from our best model (*majority rules* consensus with a correlation coefficient cutoff of 0.90) indicates that the species which our analysis identified as containing hidden genetic lineages often have ranges characterized by a larger variance in annual and seasonal precipitation, isothermality, and net primary productivity when compared to species that were not identified as containing hidden genetic lineages by our analysis (Figure 5B). And while the order of the most important traits is unstable across different models, across all models most of the traits found to be important were measurements of standard deviation (vs. measurements of mean values or life history traits) (Data S5). This suggests that the presence of variation in climate, rather than any species-specific trait or characteristic is the most identifiable driving force of within species genetic diversity for salamanders at this scale. Species traits were not a predictor of intraspecific genetic diversity in amphibians (Barrow *et al*., 2021; Amador **et al.*,* 2023) using a different measure of genetic variation within species (nucleotide diversity). Using similar methods, our results in salamanders differ from that found in mammals, where body size and range size were the most important predictors (Parsons *et al*., 2022).

These findings are somewhat consistent with other studies of salamander diversification. Reproductive mode (larval stages, direct development) and habitat (combinations of terrestrial, aquatic, arboreal) vary across species and have evolved multiple times but have not been found to directly correlate with speciation, though being a direct developer might increase diversification rates (Liedtke *et al.*, 2022). Alternatively, in one species which has intraspecific variation in habit, *Salamandra salamandra*, terrestrial-breeding individuals exhibited greater geographic genetic differentiation (Lourenço *et al.*, 2019). Not surprisingly, this species showed conflicting results in our delimitation analyses. In vertebrate clades, terrestrial organisms tend to have higher diversification rates than aquatic organisms (Wiens, 2015), but we did not have a large number of fully terrestrial species in our dataset, which might have limited our ability to detect this as an important predictor. Given that salamanders are relatively constrained in body form and ecological niches, variation in climatic variables seems like a reasonable explanation for species containing cryptic diversity. This follows the suggestion that change in climatic niche variables increases diversification rates in plethodontid salamanders (Kozak & Wiens, 2010). Diversification rates in frogs and salamanders have been shown to be higher near the tropics (Wiens 2007), so one might expect latitude to be an important predictor. However, latitude was not included in the list of predictor variables that were likely to be important (Figure 4).

### Predictive modeling as a tool to address the Linnean shortfall

Recently, Parsons *et al*. (2022) used publicly available genetic barcoding data to develop a predictive framework to identify mammalian clades most likely to contain hidden species and determine specific trait complexes that indicate where hidden mammal diversity is likely to exist. We adopted a similar approach to evaluate genetic lineages in the clade Caudata, a group which differs from mammals in several key aspects, including species richness and sampling intensity. We focused on a lower taxonomic level so there are fewer recognized species of salamanders (<1000; ‘AmphibiaWeb’, 2023) compared to the mammal dataset, making the ability to produce robust predictive models more challenging. Additionally, there was a smaller proportion of available data for salamanders than mammals (∼10% compared to 60% of described species). However, these smaller datasets might be more realistic in that they are more representative of the type of data most likely to be available for the taxonomic groups that are in greatest need of attention from taxonomists.

While the predictive models generated in this study actually have a higher overall accuracy than those used in Parsons *et al.* (2022) (see Table 3), relying on this metric alone to evaluate the performance of predictive models can be misleading (Provost *et al.*, 1998). For classification models, model accuracy depends on how well the predicted classifications match the observed classifications. While seemingly straightforward, accuracy does not account for other model characteristics that may be influencing model behavior, such as the class frequencies of the underlying dataset (Kuhn & Johnson, 2013). In cases where one class occurs at a much higher frequency than the other, a predictive model can attain a high accuracy by simply always predicting the higher class. Therefore, an important benchmark to consider when interpreting overall model accuracy is the frequency at which the majority class occurs, the no information rate (NIR). If a model’s accuracy is not significantly higher than the NIR (i.e., p-value [Accuracy > NIR]), it can remain unclear whether the model is making meaningful decisions. In our models, the overall accuracy was found to be high, but the 95% confidence intervals for the accuracy values are very wide for most of the models. In addition, because the dataset is skewed towards species classified as containing hidden diversity, the p-value [Accuracy > NIR] was found to be significant in only one model. This is important to point out because even though there are large datasets available, choosing the right analytical tools can remain challenging depending on the use of the predictive models. Beyond analytical tools, it’s also important to consider your dataset, and how the characteristics of your dataset are affecting the results you obtain. Considering the scale of not only the dataset, but also the analytical methods used and the pattern one is attempting to examine is especially important in meta-analyses, as different patterns emerge at different scales (Gurevitch *et al.*, 2018).

**Table 3.**
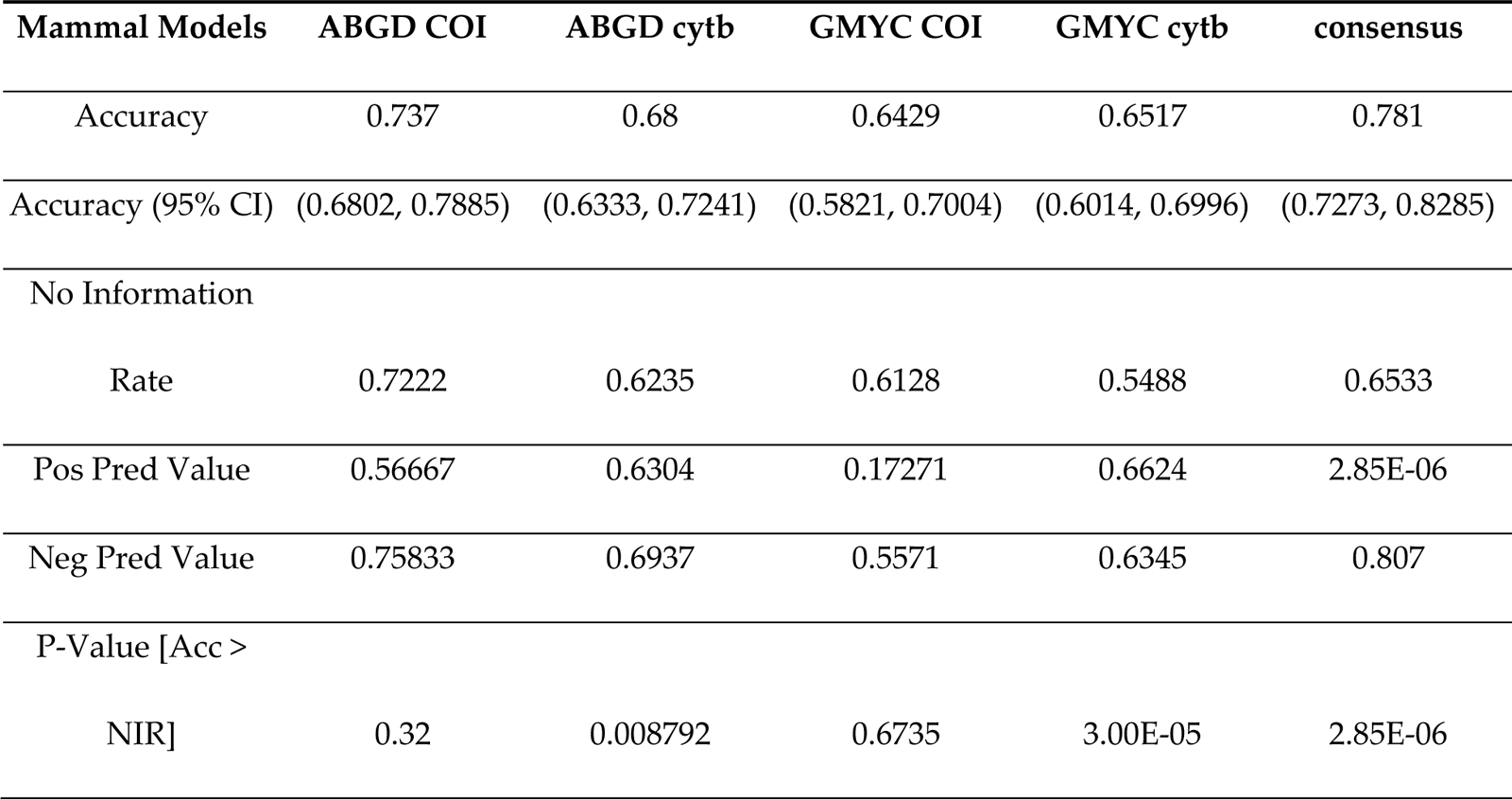
Summary of results of mammal predictive models presented in Parsons *et al*. (Parsons *et al.*, 2022). Model metrics for each random forest predictive model generated using data from the class Mammalia are shown.

## Conclusions

Here, we chose to utilize biodiversity data from *phylogatR* (i.e., genetic data for which directly associated specimen locality information is available) to avoid potential discrepancies between the distribution of the genetic and geographic data analyzed. By doing so we hoped to gain a more fine-grain understanding of how species genetic diversity is influenced by geographic and environmental factors (Leigh *et al.*, 2021). However, making this choice significantly decreased the amount of data available and led to a greatly reduced dataset. Our study included 1676 DNA barcoding sequences from the genes *COI* and *cytb* (768 and 908 sequences each, respectively). However, a 3/31/23 search of GenBank for salamander barcoding sequences from the genes *COI* and *cytb* returned a total of 17097 sequences (4468 and 12629 sequences each, respectively; see Data S6). Similarly, while we were able to obtain 1765 occurrence records tied to the genetic sequences used in this study, a GBIF search for geographic occurrences tied to salamander preserved specimens and material samples returned 675243 records (see Data S6). This study highlights the lack of genetic data with easily-associated geographic information.

The numerous benefits of making biological data more broadly available have been repeatedly demonstrated (Wüest *et al.*, 2020). And recent years have seen a significant increase in the amount of available specimen and biodiversity data. The utility of these data to address large scale patterns of biodiversity, such as those examined in this study, is enhanced by our ability to integrate and synthesize data across different data sources, types, and taxonomic groups (Heberling *et al.*, 2021). Our study highlights the importance of not just making these data available, but making them available in a way that is standardized and will facilitate integration and re-use for future generations to come (e.g., Colella *et al.*, 2021; Hardisty *et al.*, 2022).

## Acknowledgments

We thank Radford University Office of Undergraduate Research for supporting AEG through the Accelerated Research Opportunities and Research Rookies programs. We thank members of the Carstens lab for their comments on a draft of this manuscript. Funding for DJP was provided by a Presidential Fellowship from The Ohio State University and by NSF-1910623 awarded to BCC.

## Conflicts of Interest

The authors declare no conflict of interest.

## Funding

This research was funded the National Science Foundation via DBI-1911293 to TAP and DBI-1910623 to BCC.

## Data Availability Statement

Data are uploaded to Dryad (set to private for peer review) and reviewers can access it using the following link: https://datadryad.org/stash/share/4tXXsub0cPan4BqKe1KS6l5SRGwf69F5p2zOmv7QYes.

When the data is made public, the final DOI will be https://doi.org/10.5061/dryad.m63xsj474. Code related to this manuscript, including data cleaning, imputation, predictive modeling, and significance testing has been deposited in GitHub (https://github.com/parsons463/HiddenSalamanders). All remaining data are available in the manuscript and/or supporting information.

## Notes

https://doi.org/10.5061/dryad.m63xsj474

### Competing Interest Statement

The authors have declared no competing interest.

