## Supplementary material for "Predicting genetic biodiversity in salamanders using geographic, climatic, and life history traits": File S1

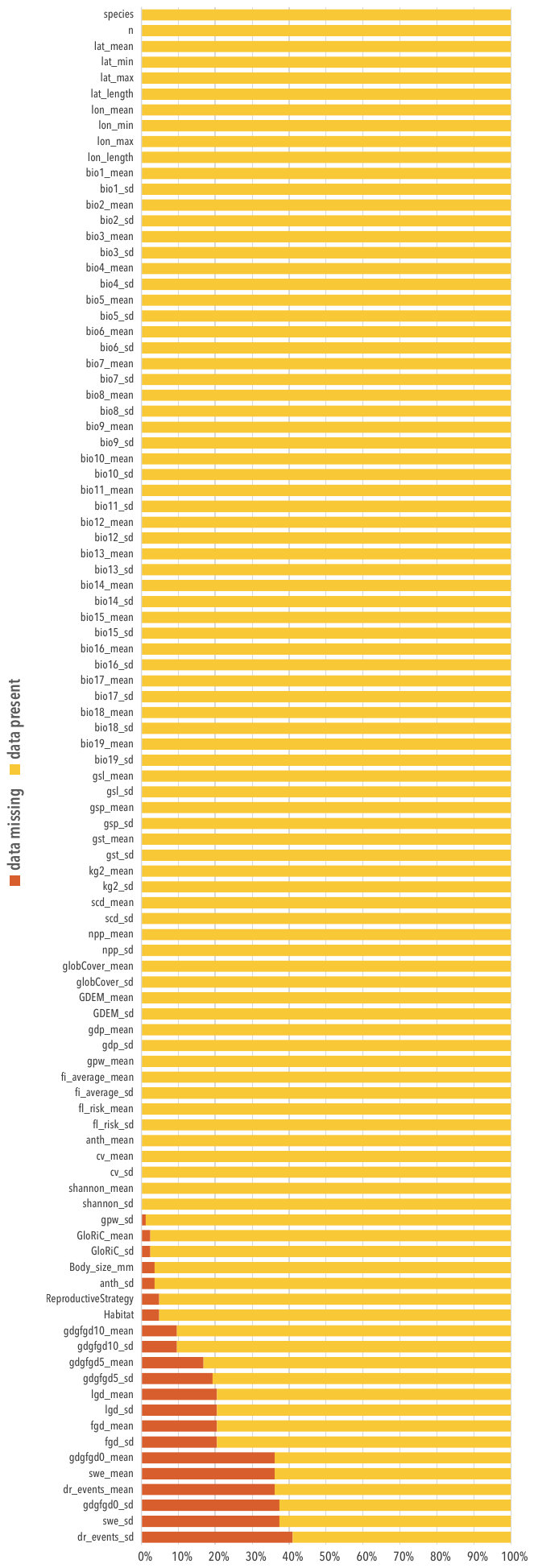

**Figure S1:** Distribution of missing data in the salamander trait database. Each bar represents a predictor variable contained within the dataset. Missing data is shown in red while previously exsisting data is shown in yellow.

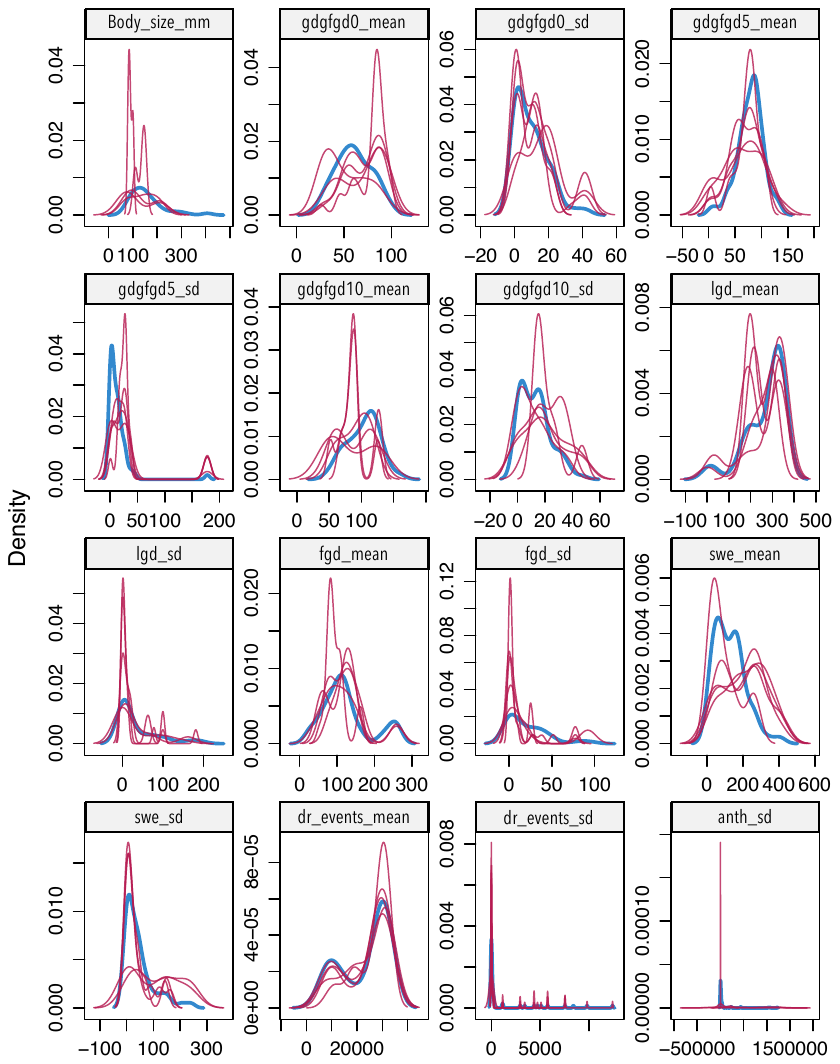

**Figure S2:** Distribution of imputed trait data. Each density plot compares the distribution of the original data (blue line) to the imputed data (red lines) for a predictor variables imputed across 15 interations. The imputed data is distributed in roughly the same shape as the original data, suggesting that the imputation process generated plausible values for missing data.

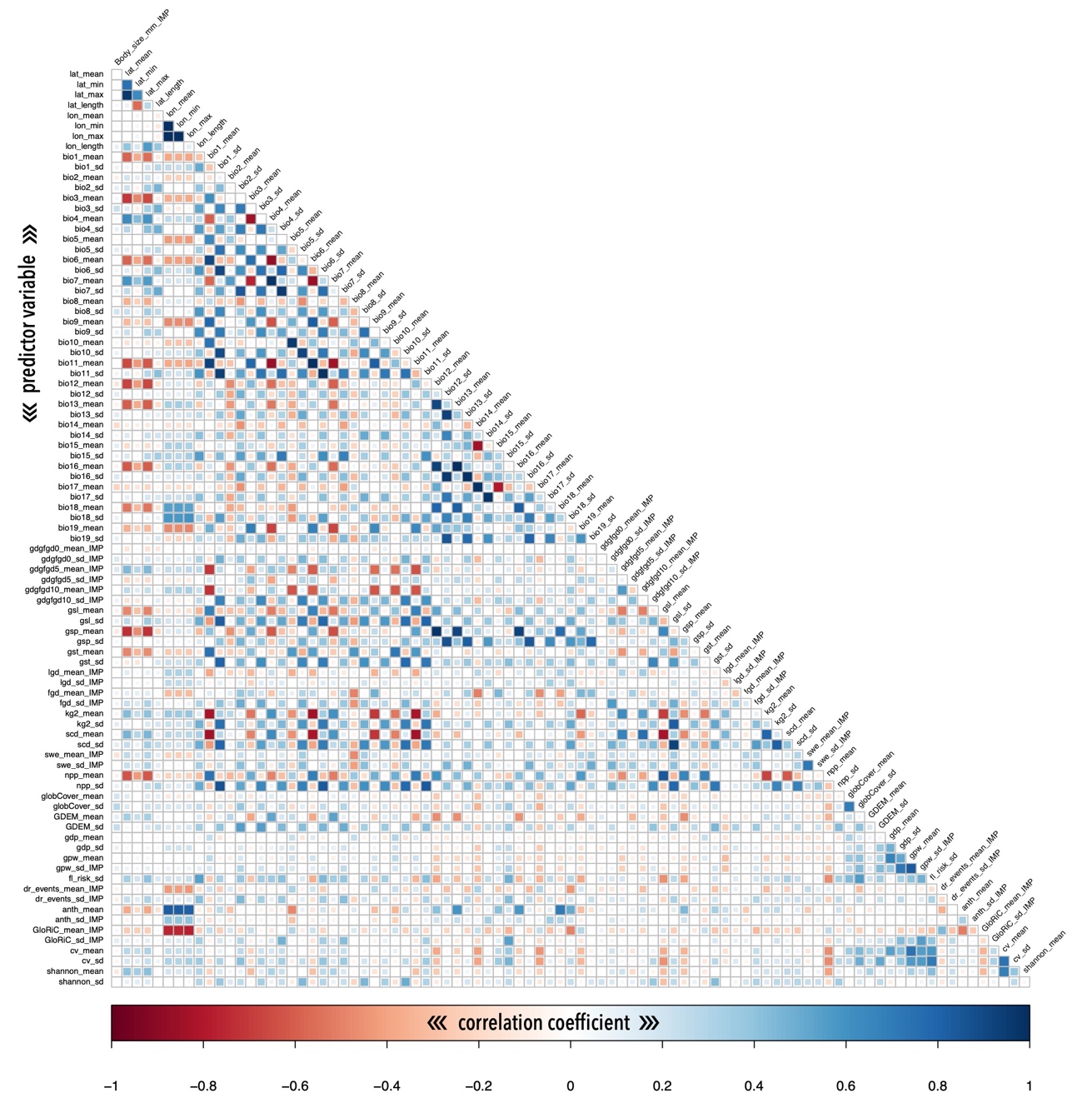

**Figure S3:** Correlation matrix of predictor variables. Correlation matrix of the predictor variables used in the final random forest analyses.

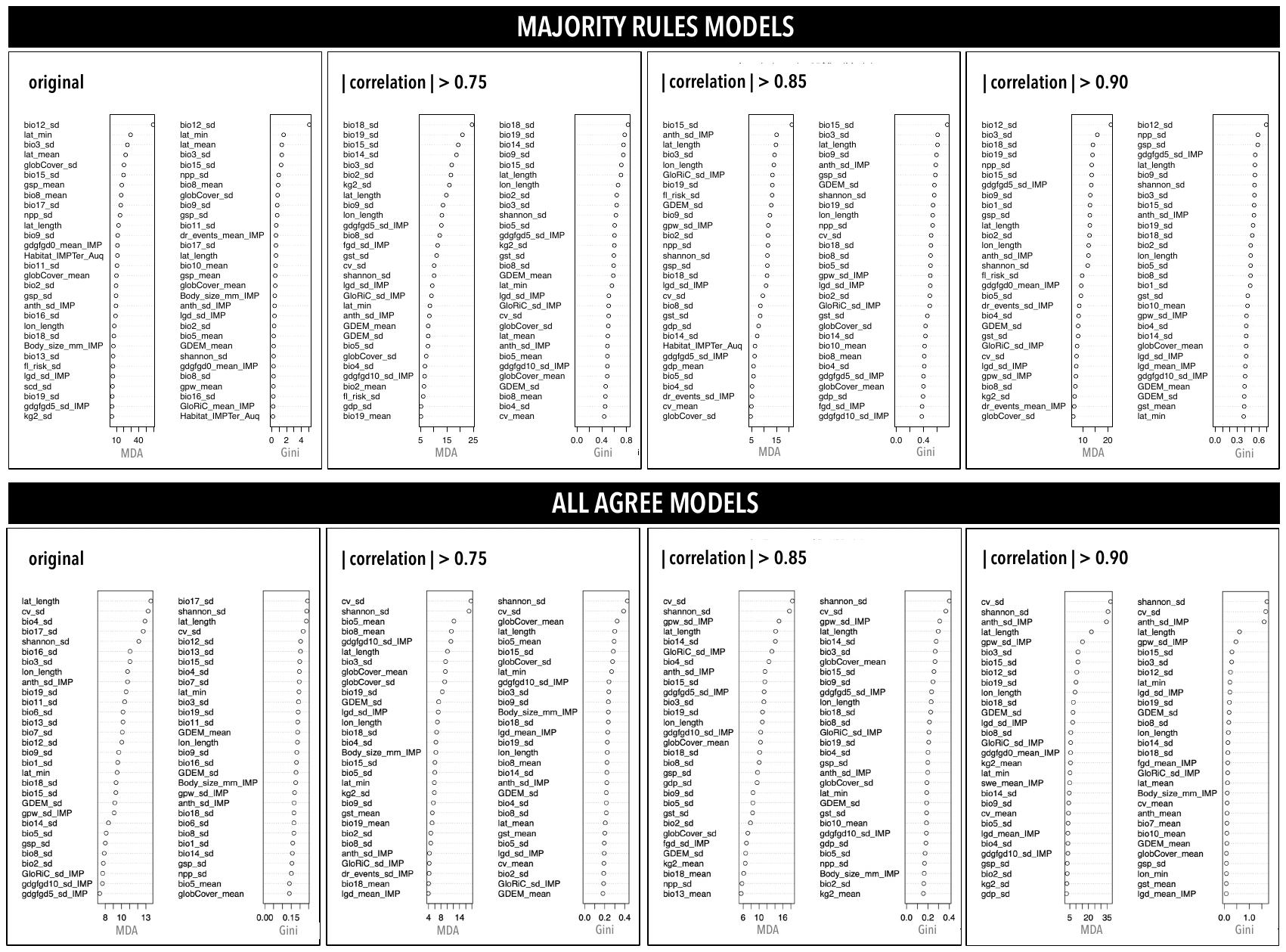

**Figure S4:** Variable importance for predictive models. Variable importance plots for all majority rules (top) and all agree (bottom) consesus models. In each plot, variable importance is ranked by mean decrease in accuracy (MDA) and Gini impurity (Gini).

| Clade | Model | Accuracy (95% CI) Length |
| --- | --- | --- |
| Caudata | MR original | 0.3694 |
|  | MR \|Correlation\| > 0.75 | 0.3694 |
|  | MR \| Correlation \| > 0.85 | 0. 3694 |
|  | MR \| Correlation \| > 0.90 | 0.3264 |
|  | AA Original | 0.388 |
|  | AA \|Correlation\| > 0.75 | 0.3178 |
|  | AA \| Correlation \| > 0.85 | 0.388 |
|  | AA \| Correlation \| > 0.90 | 0.388 |
| Mammalia | ABGD COI | 0.1083 |
|  | ABGD cytb | 0.0908 |
|  | GMYC COI | 0.1183 |
|  | GMYC cytb | 0.0982 |
|  | Consensus | 0.1012 |

**Table S1.** Comparison of model accuracy confidence intervals between salamander and mammal predictive models. In the clade Caudata, MR indicates majority rules models and AA indicates all agree models. Mammal data is summarized from Parsons et al. 2022.
